# An Embeddings Fusion Approach Predicts Disease State from Microbiome Features

**DOI:** 10.64898/2026.04.20.719747

**Authors:** Camilo Valdes, Andre Goncalves, Haonan Zhu, Boya Zhang, Hiran Ranganathan, James Thissen, Jose Manuel Marti, Car Reen Kok, Nisha Mulakken, Crystal Jaing, Nicholas Be

## Abstract

**Background:** Deep neural networks are a proven technique for working with high dimensional data because of their ability to draw-out meaningful patterns and create vector representations known as “embeddings”, which make it easier to work with learning tasks on large inputs as they capture the semantics and variance of the data. Microbial community abundance profiles are well suited for an embeddings approach due to their high dimensionality, and in this work we introduce a novel approach for generating embeddings from visual representations which encode NCBI’s taxonomic tree and microbial compositions as images, enabling the creation of embeddings that capture factors such as disease status, type, and geographical location.

**Results:** We profiled 13,534 public human metagenomes spanning 85 studies, 24 disease types, 35 countries, and 31,756 microbial species using a profiling pipeline that indexes NCBI’s nucleotide database (nt) across all kingdoms of life. Our model achieves an average classification performance of 84% in distinguishing healthy and disease conditions; 87% for disease types, 99% for body sites, and 88% for geographical locations. It also achieves a 97% accuracy when performing multi-label classification of the four factors combined.

**Conclusion:** Our work highlights the use of an embeddings approach that can encode multiple features and create efficient contextualization of profiled metagenomes derived from microbiome samples. The model’s embeddings can be used to cluster existing samples based on multiple conditions and interpretations, and new embeddings can be quickly created for new samples and fitted to existing clusters to characterize them. This has practical applications for unknown, unlabeled microbiome samples.

## 1 Background

The human microbiome operates as an “invisible organ” that is critical to nearly every aspect of human health [1, 2], and the DNA, RNA, and proteins of which a human being consists are now perceived more broadly as an interacting “hologenome” [3] composed of a host and its symbiotic microbes. Analytical methods that leverage these complex interactions could potentially be used to characterize and predict disease and healthy phenotypes [4, 5, 6], or classify samples with unknown signatures for agriculture [7, 8], forensics [9, 10, 11], and biosurveillance applications [12, 13]. These benefits are limited by the extensive heterogeneity of microbiomes between locations, individuals, body sites, and disease conditions.

Metagenomics is the study of the genetic material in a microbiome [14, 15, 16, 17], and it can provide taxonomic and functional insights into the interactions between the microbial communities and their hosts [18, 19, 20, 21]. High throughput DNA sequencing is one of the tools that can be used to capture the genetic material, and cost reductions have expanded its use [22] and allowed applications such as metagenomic whole genome sequencing (mWGS) [23] to create datasets that capture host and microbial compositions [24, 25]. Studies that use mWGS can generate large amounts of data that can be difficult to analyze and interpret; however, their scope and size make them ideal for data-driven methods that can learn unique distinguishing features within a dataset that is generalizable to samples for which little is known [26, 27].

Abundance profiles are tabular reports of mWGS analyses that detail the composition of a microbiome sample and estimate the relative quantities of its constituent microbes based on sequence abundance. They summarize the counts of DNA sequencing reads that map to microbial sequences in a database [28, 29, 30]. Each profile captures a comprehensive overview of a microbial community, and in their raw form, the profiles are numerical tables and high dimensional, as they are created by profiling against extensive microbial databases that span multiple taxonomic ranks [31]. Datasets composed of metagenomic feature data are a promising source of training data for predicting health-relevant outcomes; however, such classification efforts can be impeded the sparse and heterogenous characteristics following from the biology of their corresponding specimen sets.

Modern machine learning (ML) methods have revolutionized fields like healthcare [32] and agriculture [33], but can require large amounts of data [34, 35] to be effective. In metagenomics, an increase in the quantity of mWGS data has enabled the training of large and complex models [36], but these cannot accommodate their complex and heterogeneous features [37]. New methods for representation of integrated microbiome profiles are needed to leverage the potential of emerging state-of-the-art ML techniques.

Data representations are at the heart of ML models since they determine how well a model can learn from its training data: good representations efficiently capture the most critical information, while poor representations create noisy distributions with no underlying patterns. This is especially relevant in metagenomics, since creating physiologically-relevant representations can be a challenge due to underlying functional and genetic complexities found in mWGS data[38].

In metagenomics, ML methods have traditionally used numerical data as their input [39, 40, 41, 42], but recently, transforming numerical data into images has gained popularity [43, 44, 45]. Models pre-trained on unrelated visual tasks can generalize well [45], eliminating the need for costly training runs and overcoming limitations of small datasets.

We present an ML model that uses “*visual* “ representations of microbiome abundance profiles, known as “Microbiome Maps” [43], to create embeddings that encode multiple factors. The representations are highly interpretable images that display microbial compositional patterns, and their embeddings allow interpretable characterization of human gut microbiome species, enabling an understanding of their potential roles in causing or indicating human diseases.

The model constructs a unified representation space and creates embeddings that can form clusters that enhance the delineation of biologically meaningful sub-groups and reveal patterns across multiple metadata dimensions. Our results show that computer vision models can be trained with “visual” representations of metagenomic profiles, and enable the classification of human health-relevant phenotypes with high accuracy and strong potential for future generalizability.

## 2 Materials and Methods

We use a custom bioinformatics pipeline to profile 13,534 public metagenomes, spanning 85 studies, two conditions, 24 disease types, 35 regions, four body sites, and 31K microbial species. [46]. The pipeline uses NCBI’s *nt* database to profile against all kingdoms of NCBI’s taxonomy (“Bacteria”, “Viruses”, “Eukaryotes”, “Archaea”) using the Centrifuge and Recentrifuge classifiers [47, 48] to create tabular reports of taxonomic counts of detected microbes. Profiles are also curated with their clinical metadata.

Data are normalized by applying a centered log-ratio (CLR) transformation to the abundance counts [49]. CLR is a reference-based normalization method commonly used in compositional data analysis (CoDA [50]), in which data represents parts of a whole. Compositional data cannot be analyzed effectively using standard statistical methods due to the constraint that the sum of all components (i.e., taxon abundances) equals a constant (1 or 100%). CLR alleviates this by scaling each component relative to the geometric mean and provides a central value that reflects the overall scale of the data. Recently, it has been proposed that proportion-based normalization methods perform better than CLR-like methods in ML tasks [42], but only in amplicon sequence variant (ASV) data, which does not have the taxonomic resolution of mWGS.

**Table 1:** Number of samples in each disease category and condition. Samples are annotated with “control” or “disease” labels, but some datasets like the Human Microbiome Project’s Irritable Bowel Syndrome (IBS) do not have a disease label.

| Disease Type | Samples | No. Control | No. Diseased |
| --- | --- | --- | --- |
| Autoimmune | 306 | 101 | 205 |
| Cancer | 1,812 | 967 | 845 |
| Cardiovascular | 930 | 335 | 595 |
| Cirrhosis | 308 | 190 | 118 |
| Constipation | 10 | 5 | 5 |
| Dermatologic | 450 | 153 | 297 |
| Diabetes | 233 | 86 | 147 |
| Diarrhea | 399 | 226 | 173 |
| Gastrointestinal | 509 | 279 | 230 |
| Genitourinary | 20 | 8 | 12 |
| GI Neurological | 648 | 338 | 310 |
| Healthy Subject Study | 1,581 | 1,581 | 0 |
| Hormonal | 93 | 43 | 50 |
| Immune | 156 | 27 | 129 |
| IBD | 1,821 | 527 | 1,294 |
| IBS | 105 | 42 | 63 |
| Macular Degeneration | 113 | 44 | 69 |
| Metabolic | 20 | 10 | 10 |
| Musculoskeletal | 47 | 22 | 25 |
| Neurological | 2,354 | 1,074 | 1,280 |
| Oral | 191 | 78 | 113 |
| Premature Born | 418 | 4 | 414 |
| Pulmonary | 856 | 434 | 422 |
| Seafaring Syndrome | 154 | 99 | 55 |

### Microbiome Map Images

Tabular profiles are converted into images with the Jasper tool [43, 51], updated for CLR-transformed data and a color palette that supports negative and positive ranges. An image is created for each profile, with no neighborhood labels (Figure 2B), as they are only useful for inspecting single images. Pixels represent a single microbe, with color corresponding to their CLR abundance, and positions in *all* images remain *constant* (only color changes), creating learnable visual patterns that reflect a microbe’s compositional changes (Figure 2).

**Figure 1:**
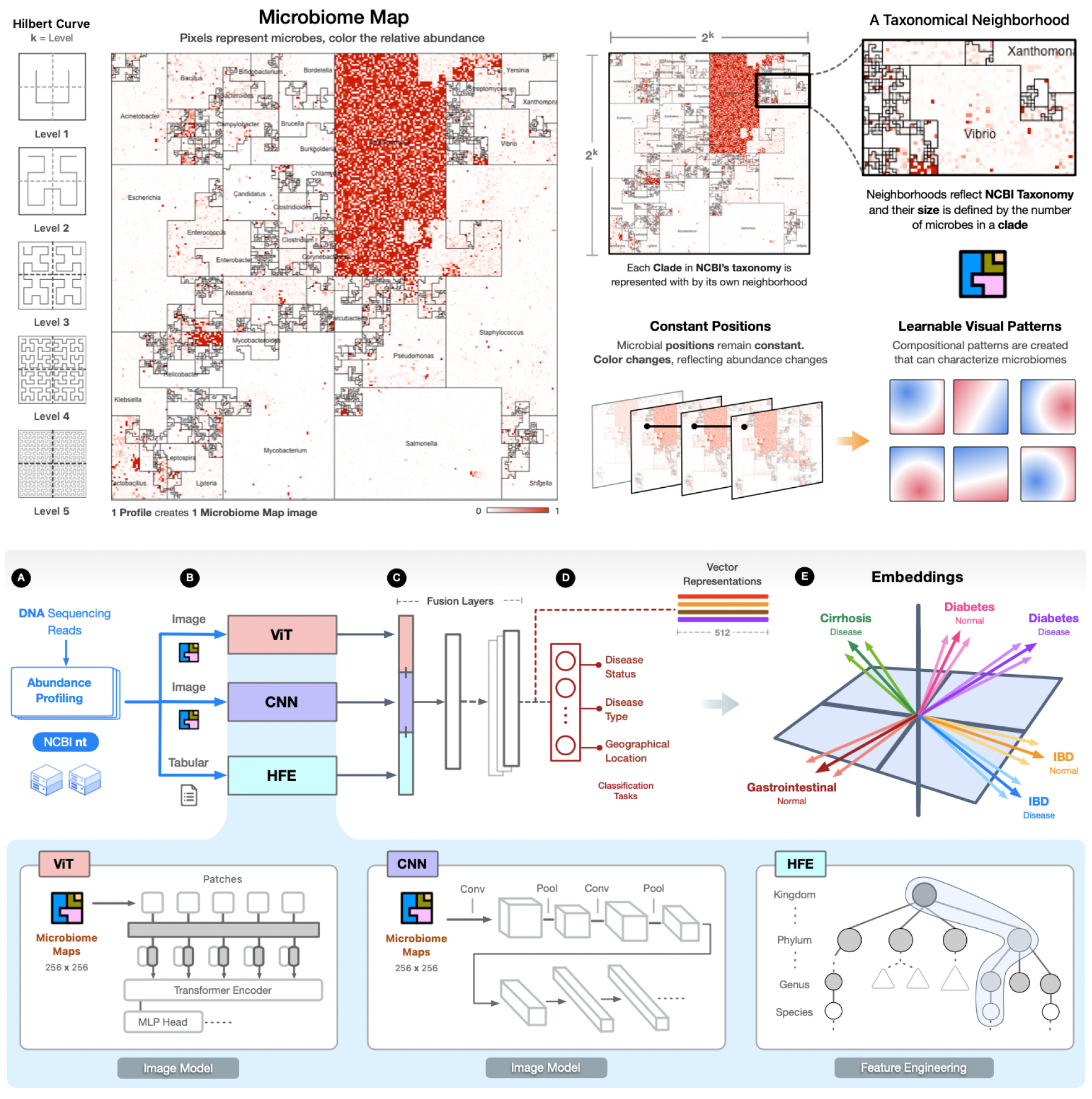
Microbiome Maps are image representations of mWGS profiles. One image corresponds to one profile; pixels represent microbes and color their relative abundance. (**A**) mWGS data are profiled against NCBI’s nt database. (**B**) HFE’s inputs are tabular profiles, ViT and CNN models use image profiles. (**C**) Feature vectors are integrated with HFE features to create a unified representation space. (**D & E**) Final latent representations are created as a unified embedding space that integrates learned features from disease status, type, body site, and geographical location.

**Figure 2:**
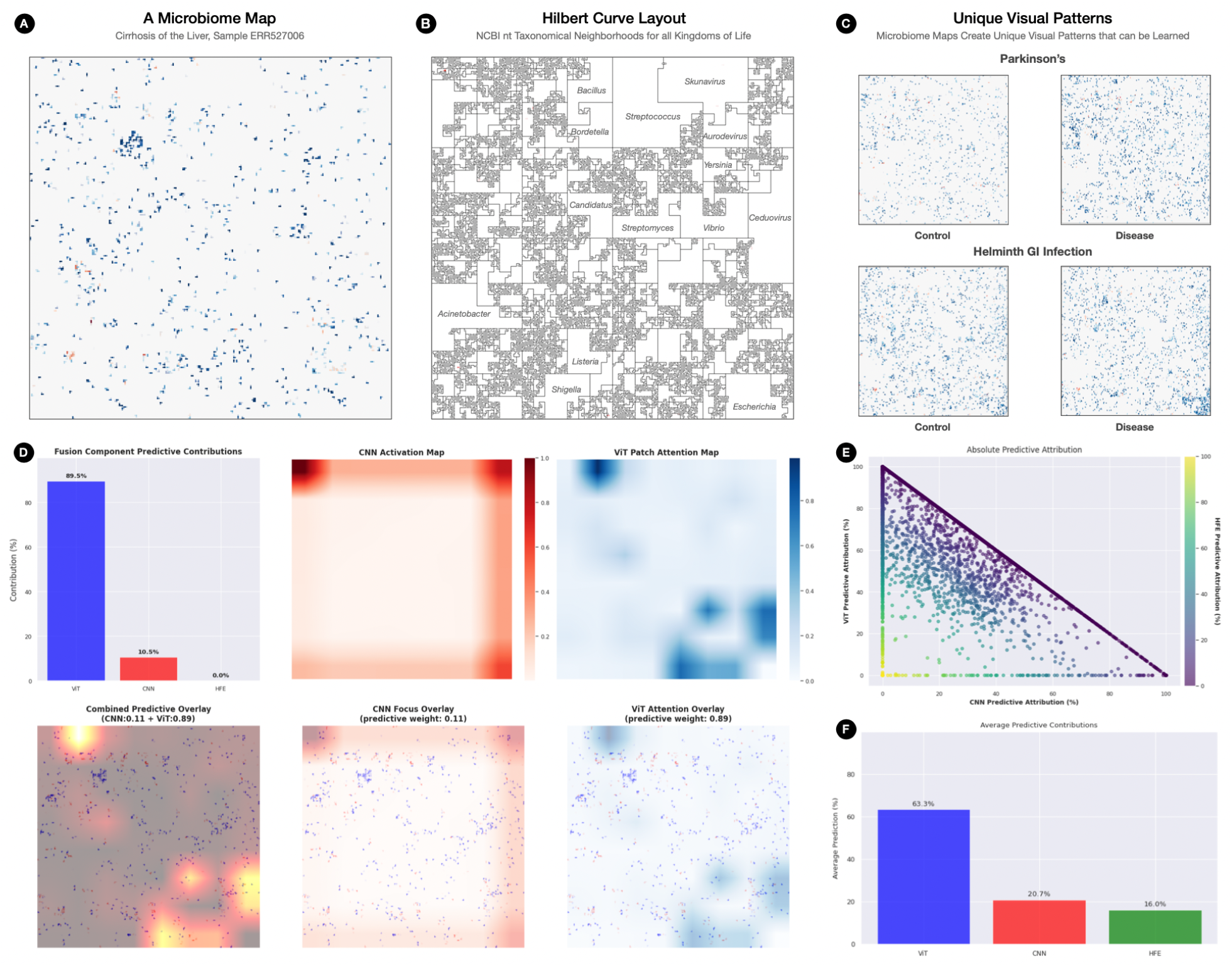
(**A**) Microbiome map representing a single abundance profile. (**B**) Hilbert curve layout of taxonomic neighborhoods and microbial positions used in *all* maps. The areas represent the taxonomic distribution of NCBI’s “nt” database at Genus and Species levels. (**C**) Four representative microbiome maps highlighting the unique visual patterns each disease creates. (**D, E**) Attention and activation maps highlight the areas that the ViT and CNN models focus on. (**F**) Fusion model heatmaps combine the CNN’s and ViT’s focal maps, weighted by the fusion head, to show the regions that most influenced the final classification.

### Hierarchical Feature Engineering

We also leverage NCBI’s taxonomy by using hierarchical feature engineering (HFE) [52] to create features at different lineages and taxonomic paths. These features represent the most informative taxonomic elements and aggregated abundances, which capture phylogenetic compositions and their relationships to disease variables.

### 2.1 A Fusion Model for Metagenomics

Late fusion is an ML technique where models are trained separately on the same data and their high-level features combined to make a decision [53, 54], and contrasts with methods in which features are combined *before* training [55]. An advantage is that models can be trained independently with different optimization and featurization schemes. A disadvantage is that it increases model complexity and training times.

Vision Transformers (ViT) [56] are neural architectures that use transformers [57] in vision tasks. They divide an image into a grid of smaller patches, flatten them into a 1D vector, and project them into a higher-dimensional space using linear transformations (Figure 3). “Self-attention” [57] is computed on these vectors by calculating scores between patches, and used to weight each patch’s contribution to the representation, enabling the model to focus on relationships across different regions of an image.

**Figure 3:**
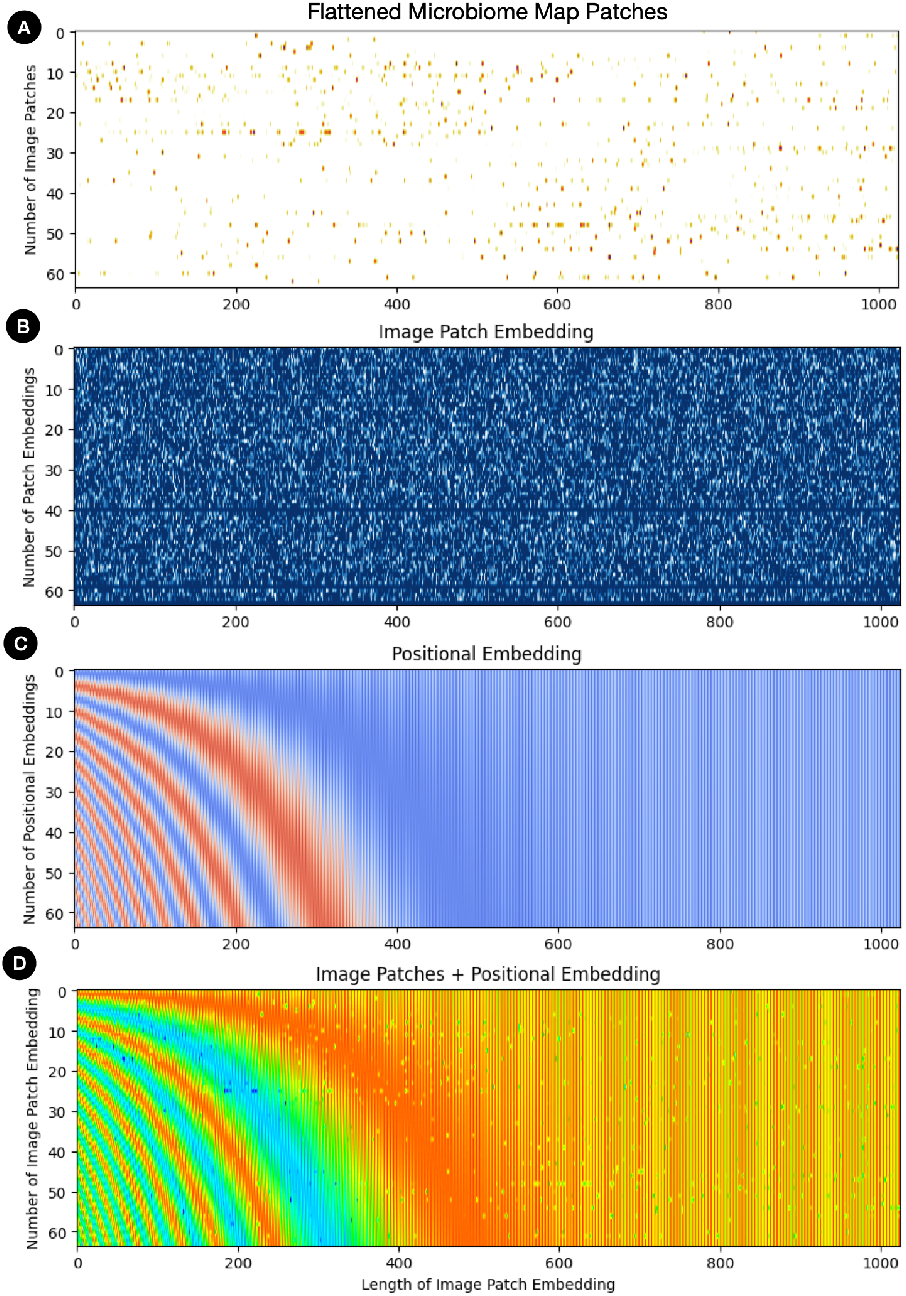
ViT. (**A**) Maps are split into patches and flattened. (**B**) Patches are transformed into vectors which can be uniquely identified (horizontal lines). (**C**) Positional embeddings retain information about the position of each patch. (**D**) Patch and positional embeddings are combined to create a unique representation. Visualization as in [61].

Our ViT implementation follows standard architectures and includes image tokenization and positional embeddings (Figure 3), followed by four transformer blocks [56] that allow the model to integrate the spatially discrete abundance patches within a map. The blocks split the input (patch-level feature vectors) into subspaces (i.e., distinct projections that let each head “see” different feature combinations) so the model can learn a variety of disease patterns in parallel. Our ViT model contains 10.7 million trainable parameters, and outputs a feature vector of size 512.

Convolutional neural networks (CNN) [58, 59] are another type of architecture for working with images. They work by applying convolutional filters to small local patches (e.g., 3×3 pixels) to identify features like edges, textures, and shapes. By stacking convolutional layers with pooling layers that summarize features, CNNs build hierarchical feature representations and can learn to recognize complex visual patterns with high accuracy and speed [60]. We implement a custom sparse-aware CNN that consists of five convolutional blocks, with spatial attention masks applied after the second and fifth blocks to suppress low-activation background areas and focus the model’s feature extraction on regions of stronger activation (taxonomical neighborhoods, i.e., “hot spots” of very abundant microbes). Each block consists of convolution, BatchNorm, ReLU, dropout, and max-pooling stages. Activation masks are applied at two points in the network, after the second block and after the fifth block. After masking, the feature maps are flattened and passed through a two-layer classifier that outputs either probabilities for multi-label tasks or raw logits otherwise. This architecture preserves efficiency while improving focus on taxonomical neighborhoods.

#### Fusion Architecture

Our fusion model combines a CNN and ViT to learn the unique visual patterns in our representations (Figure 2, panel “C”). This leverages the CNN’s ability to learn *local* patterns with the ViT’s strength in modeling *global* context, making it well-suited for the spatial details of the maps.

The model builds a combined feature vector **f**_concat_ (equation 1) by concatenating the CNN output **f**_CNN_ (512), the ViT output **f**_ViT_ (512), and the HFE output **f**_HFE_ (512) and passing it through two fully connected (FC) linear layers. In the first FC layer, **h**_1_ (equation 2), **f**_concat_ is projected into a lower-dimensional latent space with an activation that injects non-linearity and reduces dimensionality. **h**_1_’s output is passed to the second FC layer, **h**_2_ (equation 3), which further refines the latent representation from **h**_1_ to **h**_2_.

These layers provide dimensionality reduction on **f**_concat_ and transform it to a 512-dimensional embedding, **o** (equation 4), whose output is constrained to [0, 1], representing the probabilities of *C* classes. Equation 6 then creates a unified representation that integrates complementary local, global, and HFE signals.

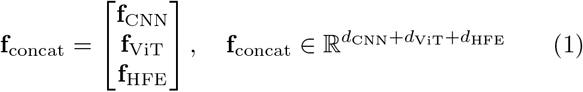

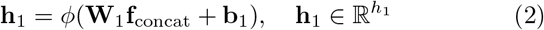

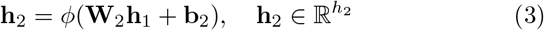

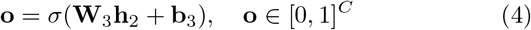

### 2.2 Embeddings for Abundance Profiles

Embeddings are vectors of numbers in continuous low-dimensional space, useful for capturing the variance, structure, and features of data. In visual tasks, they are used to create vector representations of images and ideal for clustering and classification [62, 63].

In metagenomics, embeddings have been used to encode protein information from ontology databases like UniProt and SwissProt [64]. These approaches are relatively efficient, providing a way of encoding key functional features without the need of sequence-based analysis methods [65]. They are not perfect however, as sequencing data still has to be processed (which can be computationally expensive).

We generate our embeddings using the fusion model trained with a multi-label classification scheme, making a forward pass with each map to extract **e** (Equation 5) as the abundance profile embedding. The dimension of **e** was chosen empirically as 512, and is a hyperparameter. This size provided a good tradeoff in representing the visual patterns in our maps and generated the best performance in clustering and generalization evaluations (see supplementary materials). Note that **e** is a dense representation and does not apply an activation function *σ* like **o**.

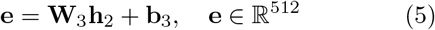

## 3 Results

We evaluate our models with 3-runs of 5-fold cross validation for “disease status” using binary classification; for multi-class classification using “disease type”, “body site”, or “geographical location”; and for multi-label classification which combines all factors.

To establish a baseline and verify whether off-the-shelf models could meet our needs, we evaluated six publicly available models from TorchVision [66], pretrained with “ImageNet 1K” [67]. We aimed to leverage transfer learning as [45] did with mass spectrometry (MS) image representations of prostate cancer biopsies. Performance results for 3 CNN models and 3 ViT models are reported in tables 4-11 of the supplementary materials. While they showed acceptable performance on some of the tasks (table 10 supplementals), their overall performance reflected a fundamental domain mismatch: ImageNet pre-training optimizes for dense, textured natural images with color gradients, while microbiome maps contain sparse, symbolic clusters representing microbial abundance.

Given their relatively low performance, we designed custom ViT and CNN architectures for microbiome maps. The models are optimized for images containing sparse, isolated signal patches against backgrounds of undetected species, with each patch’s position and intensity carrying strict taxonomic meaning rather than forming textures or shapes.

### 3.1 Model Training & Evaluation

Our data consists of 13K microbiome map images, with 90% used for training and validation (80%/20%), and 10% used for testing (holdout). We first train and evaluate our custom ViT and CNN models on the same data and then integrate them as submodel backbones into the fusion architecture by removing their classification heads and routing output features to the trainable layers *h*_1_ and *h*_2_.

For evaluating the “disease status” of control *vs*. disease samples, our CNN achieves a mean AUC of 0.78, and the ViT 0.78 (macro AUROC; for reference, the trainable and frozen fusion models reach 0.85 and 0.82, respectively). In evaluating the 24 disease types in multi-class classification, the CNN achieves a mean AUC of 0.93, with the ViT 0.96 (the trainable variant of the fusion model achieved 0.98). For classifying the 35 locations, the CNN had a mean AUC of 0.96, and the ViT 0.98 (trainable fusion model achieved 0.99). For the four body sites, both models had a mean AUC of 0.99 (more in section 7.3 supplementals).

We also tested whether our models could predict all factors at once, and sections 7.3.3 and 7.3.4 of the supplementals reports two ways to measure success: *per-label* and *per-image* accuracy. *Per-label* accuracy is the average prediction rate across each individual label; *per-image* accuracy requires every label for a given sample to be predicted correctly, and any single misclassification renders an incorrect prediction. Depending on the application at hand, misclassification in totality may not be a relevant metric at the application level, as certain categorical labels may not be critical, for instance, in a clinical environment.

Results from these early training demonstrations indicate the promise of implementing this approach to predict health state via a fusion model (AUC 0.84) for a wide range of disease conditions. While it may be readily achievable to achieve comparable predictive performance when isolating a given analysis to a single disease condition, the capacity demonstrated here, to apply a single fusion model for prediction of multiple disease states from one consistent microbiome feature set, represents a substantial additional contribution, given the wide range of underlying mechanisms that may be specific to different disease etiologies.

Finally the latter demonstration of extremely high level of accuracy in body site prediction reflects the anticipated high degree of compositional differences between microbiomes derived from distinct locations across the human body.

#### 3.1.1 Fusion Model Performance

We tested two training setups for the fusion model: (1) freezing the submodels after training them independently, and (2) fine-tune the submodels together with the fusion layers. While freezing the submodels preserved each model’s specialized feature extraction capabilities and reduced training complexity, we found that end-to-end fine-tuning of the entire fusion architecture delivered superior performance across most metrics (Table 2 and Figure 4).

**Table 2:**
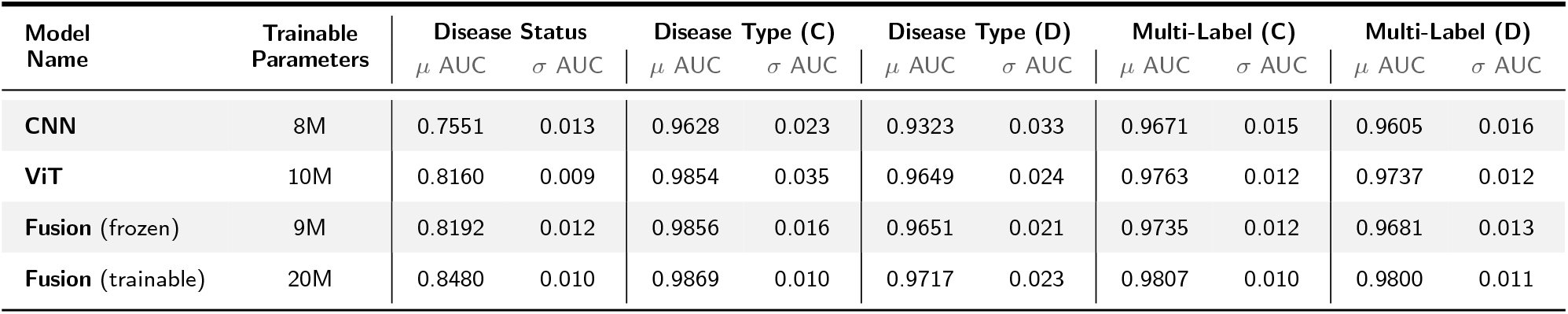
Holdout set AUC metrics (*N* = 1, 350). Our custom models are trained with our set of 13K microbiome map images. The “Disease Type” trial tested a multi-class task with control samples (C), and disease-only samples (D). The multi-label task also included trials with control samples (C) and disease-only samples (D). Control samples are included to set a disease baseline.

| Model Name | Trainable Parameters | Disease Status |  | Disease Type (C) |  | Disease Type (D) |  | Multi-Label (C) |  | Multi-Label (D) |  |
| --- | --- | --- | --- | --- | --- | --- | --- | --- | --- | --- | --- |
| | | $\mu$ AUC | $\sigma$ AUC | $\mu$ AUC | $\sigma$ AUC | $\mu$ AUC | $\sigma$ AUC | $\mu$ AUC | $\sigma$ AUC | $\mu$ AUC | $\sigma$ AUC |
| CNN | 8M | 0.7551 | 0.013 | 0.9628 | 0.023 | 0.9323 | 0.033 | 0.9671 | 0.015 | 0.9605 | 0.016 |
| ViT | 10M | 0.8160 | 0.009 | 0.9854 | 0.035 | 0.9649 | 0.024 | 0.9763 | 0.012 | 0.9737 | 0.012 |
| Fusion (frozen) | 9M | 0.8192 | 0.012 | 0.9856 | 0.016 | 0.9651 | 0.021 | 0.9735 | 0.012 | 0.9681 | 0.013 |
| Fusion (trainable) | 20M | 0.8480 | 0.010 | 0.9869 | 0.010 | 0.9717 | 0.023 | 0.9807 | 0.010 | 0.9800 | 0.011 |

**Figure 4:**
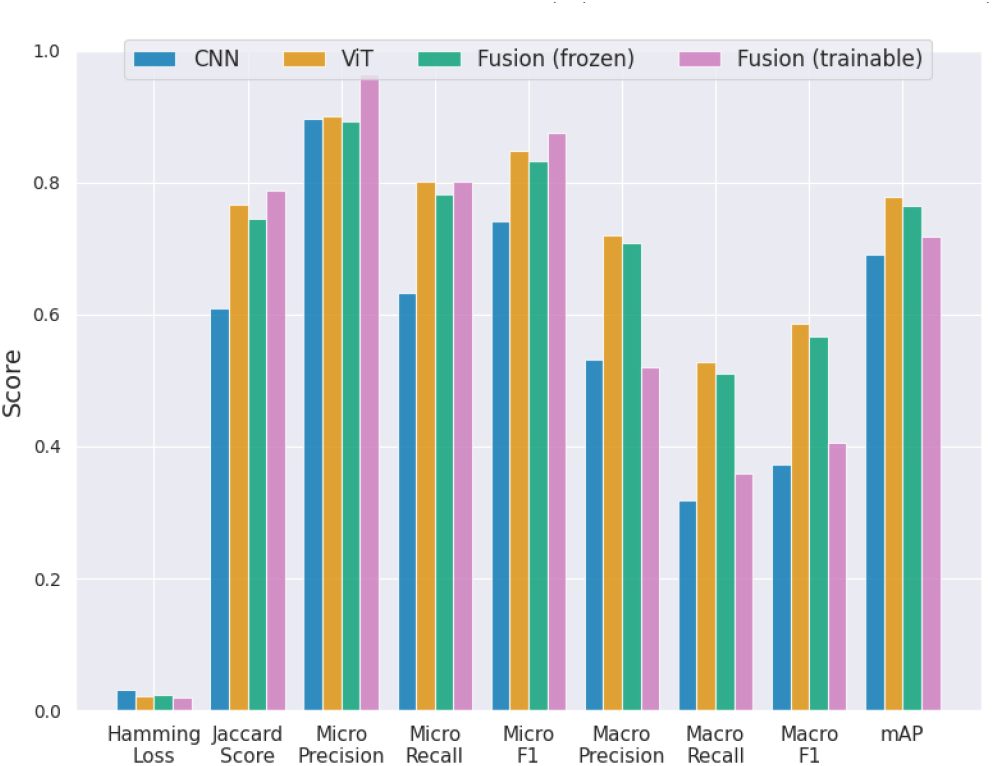
Multi-label (w/ controls) metrics for CNN, ViT and Fusion models. Trainable and frozen fusion models represent models for which their corresponding backbones are trainable or not.

**Figure 5:**
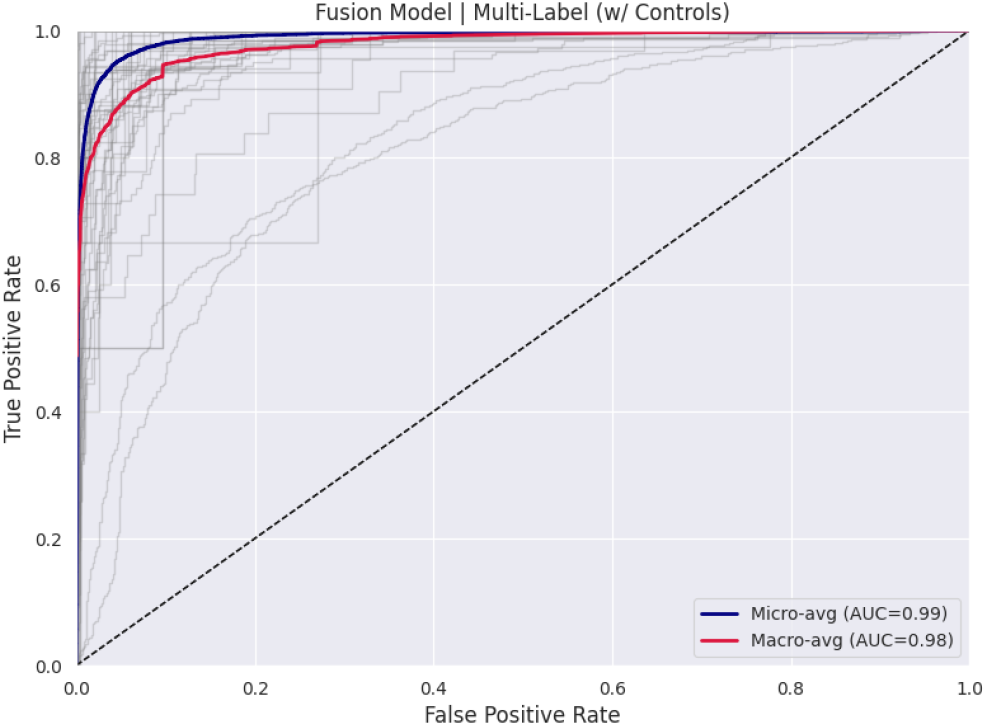
Trainable fusion model micro AUC above macro AUC shows minimal class imbalance effects in multi-label (D) trial. Micro is frequency-weighted across labels, macro is unweighted mean.

The fusion model architecture (frozen) achieved consistently strong overall performance in the multi-label task with low Hamming loss (0.022) and high micro-precision (0.893). Micro-macro gaps (precision: 0.893 vs 0.707; recall: 0.782 vs 0.511) show significantly improved class imbalance handling across labels. A strong Jaccard score (0.745) and mAP (0.764) show complementary feature fusion, with balanced recall metrics and improved micro-recall (0.782) showing a better precision-recall compromise.

The trainable fusion model delivered the best overall performance in the multi-label task with the lowest Hamming loss (0.019) and highest micro-precision (0.965), indicating the most accurate predictions. The micro-macro gaps (precision: 0.965 vs 0.519; recall: 0.801 vs 0.358) show good class imbalance handling and consistent label assignments. The highest Jaccard score (0.787) and third highest mAP (0.718) suggests strong performance on frequent labels but struggling on rare ones. The full table of results for each of the seven tasks can be found in tables 5-11 of the supplementary materials.

We evaluated our models with seven tasks, and the fusion architecture was the best overall on six, with the trainable variant the best on disease status, disease type (with controls), disease type (disease-only), country, and multi-label (with controls) based on accuracy, Hamming loss, Jaccard, and micro-F1. The frozen variant was best on body site, and our custom ViT was best on multi-label (disease-only), where it had the highest accuracy, Jaccard, and micro/macro-F1.

Our performance observations indicate that the fusion model architecture confers benefits beyond the individual sub-approaches separately. Even in cases where there is only marginal such improvement, this approach will likely provide for improved flexibility of the fusion model to generalize, which we explore further in the following section.

### 3.2 Embeddings for Unseen Microbiomes

To evaluate the fusion model’s generalization capability for microbiomes not previously seen by the model, we use leave-one-out (LOO) cross validation with 85 studies: one study is left out completely from training, and the models are trained with the remaining ones. Ideally, we would train the model *k* times (where *k* = 1, 2, 3, …, 85), but some studies contained a small number of samples, and we exclude them such that |𝒟_*k*_|> 200 for all *k* ∈ {1, 2, …, 85}, where |𝒟_*k*_| is the number of samples in the *k*-th study (section 7.4 supplementals).

We also evaluate the structure of the model’s representation space in the context of disease clusters, and to do so we construct clusters based on sample embeddings and assess their alignment with expected distributions using Pearson’s *χ*^2^ goodness-of-fit test. Specifically, we evaluate the *p*-value associated with the test statistic, *χ*^2^, under the null hypothesis using a significance level *α*. This approach quantitatively evaluates whether the model’s embedding-derived clusters encode the expected *visual* patterns created by microbial “hot spots” in samples from unseen datasets.

In this analysis, new embeddings are created from microbiome maps derived from studies unseen by the model during training, and we let the null hypothesis *H*_0_ be: “The new embeddings generated by the fusion model follow the expected cluster distribution”, and the alternative hypothesis *H*_*a*_: “The new embeddings do not follow the expected distribution”. We perform the *χ*^2^ test for new embeddings (section 7.4 of supplementals), and compute the *p*-value corresponding to the test statistic using the chi-squared distribution with *k* − 1 degrees of freedom, such that if *p* ≤ *α*, we reject *H*_0_ (with *α* = 0.05), and if *p > α*, then we fail to reject *H*_0_.

We observed no significant differences between new embeddings and existing disease clusters: we were unable to reject the null hypothesis, *H*_0_, for any of the studies. This indicates that new embeddings for unknown microbiomes accurately reflect the expected clustering patterns, and they should generalize to previously unseen data. This suggests that our model is capable of capturing the underlying metagenomic and abundance patterns and bodes well for its performance in real-world health-based applications where mWGS data are continuously produced and evaluation by a generalizable model that has not previously seen the emerging data is a requirement.

## 4 Discussion

Metagenomic datasets derived from the human microbiome represent a rich feature set for characterizing human disease. The ability of these features to represent physiological parameters and train ML models for predicting physiological responses and categorical clinical data has been shown in numerous targeted instances [68, 69, 70]. Challenges remain, however, in representing such datasets such that they may be robustly and accurately assessed by emerging ML methods, as microbiome data are particularly sparse compositional, and high-dimensional (*p* ≫ *n*) [71]. Our efforts demonstrate that an image-based fusion ML approach can be implemented to improve microbiome classification.

Inter-individual and inter-cohort heterogeneity [72] represent a substantial challenge in evaluating the accuracy of ML models built from microbiome datasets. Our multi-label evaluation scheme measures the ability to classify disease status, type, body site, and location simultaneously from a single microbiome map image. Unlike the individual classification tasks, this approach treats all biological and geographical factors as interconnected dimensions that must be predicted together, reflecting the reality that microbiome patterns are influenced by a broad range of demographic factors and therefore encode multiple scales of information simultaneously. This design acknowledges that a microbiome map’s spatial patterns contain layered information: the same discrete patches and intensity variations simultaneously indicate the host’s health status, the body site sampled, the specific disease present, and the population’s geographical context. An evaluation scheme with the capacity to assess multiple such variables provides flexibility for future efforts which may seek to predict a range of phenotypes and outcomes, depending on the investigation goal.

### Supplementing Tabular Data for Improved Performance

The inherent difficulties in microbiome datasets are sometimes addressed via methods to transform these data and assess differential feature abundance; however, challenges remain with analyses producing a range of outcomes depending on the method applied [73]. This suggests inherent shortcomings in such tabular datasets, which could be addressed or compensated for through integration with alternative representations, such as images [43]. Indeed, our results demonstrate improved embeddings separation performance of our fusion model relative to each tabular dataset (≈ .6 to ≈.9) (section 7.1 supplementals), indicating that multimodal models improve results for clinical prediction applications, which is consistent with observations in other domains [74].

Section 7.1 and 7.2 of the supplementary materials contain results of evaluating eight popular classifiers using the tabular CLR-transformed data. This analysis serves as a baseline to assess how well conventional classification methods perform on compositionally corrected microbial abundance data before exploring and comparing to computer vision approaches.

We selected linear methods (Logistic Regression, Linear SVM), distance-based approaches (k-Nearest Neighbors), non-linear kernel methods (RBF SVM), tree-based algorithms (Decision Tree, Random Forest), and ensemble boosting techniques (AdaBoost, Histogram Gradient Boosting) to allow for a broad and flexible assessment of the tabular CLR datasets [75, 76]. While the accuracies in supplementary figure 16 quantify how well each algorithm performed on the tabular data, we also wanted to understand how they arrive at those decisions by investigating their decision boundaries and embeddings (supplementary figures 17, 18). While these classifiers achieved accuracies with mean of 0.70, their decision boundaries and embeddings revealed a key limitation: they failed to capture meaningful biological structure in the data. Unlike our computer vision embeddings (Figure 6), the clusters formed by conventional methods do not reflect the underlying compositional relationships between microbial communities. These observations reinforced our hypothesis that tabular representations of metagenomic data may be limiting, and that an approach integrating image data modes could compensate.

**Figure 6.**
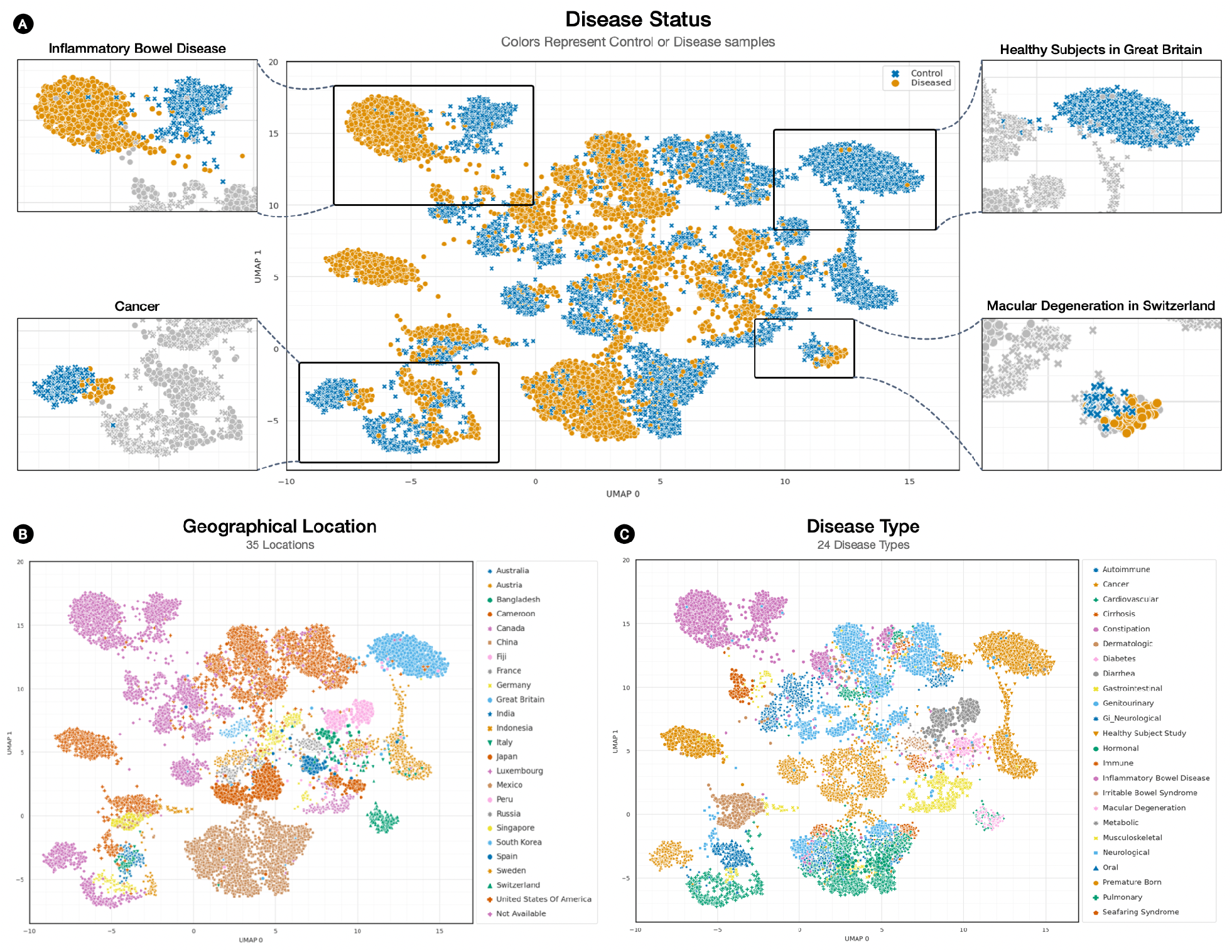
Multi-label embeddings for single metagenomic abundance profiles represented as microbiome maps. (**A**) Cluster by multi-label embedding, colored and shaped by disease status with highlight boxes displaying selected diseases. (**B**) Same as “A” panel, but now colored by geographical locations. (**C**) Same as “A” panel, but now Colored and shaped by disease type.

**Figure 7.**
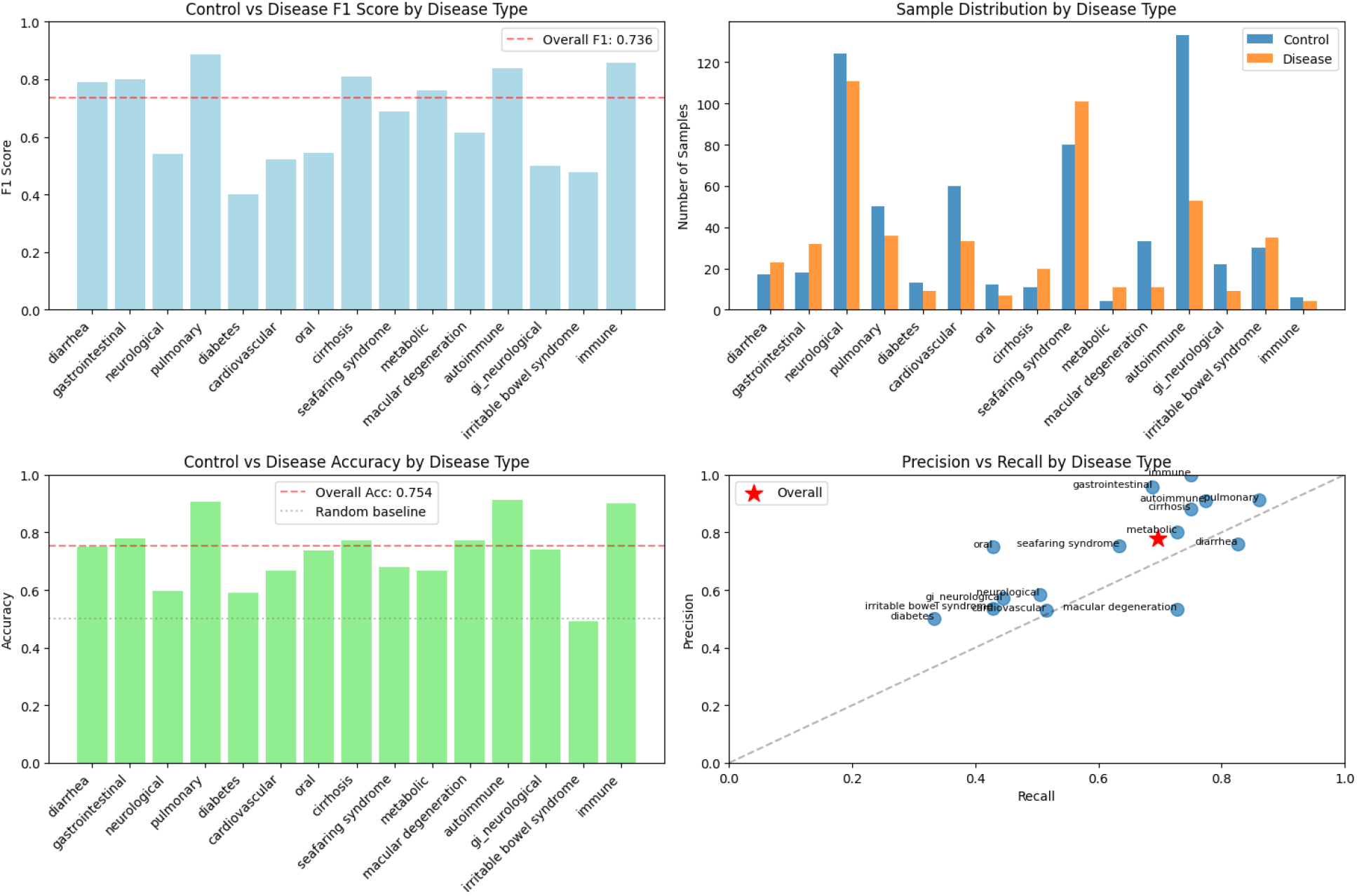
Fusion model performance on holdout set (n=1354) for disease status by disease type: F1 (top-left) and accuracy (bottom-left) bars are shown against overall baselines (dashed), with sample counts by type (top-right). Performance varies by type as many exceed the overall F1 (0.736) while a few fall below 0.6, consistent with uneven class sizes. Red star marks overall (micro) operating point (0.75 precision, 0.72 recall), indicating solid but improvable recall for some types.

### Disease State Prediction with Heterogenous Datasets

Increasing attention is being paid to the capacity of human microbiome features specifically for prediction of clinically relevant outcomes and risk stratification [77]. Such applications would provide critical information from complex metagenomic data that could supplement clinical decision support, which has been demonstrated in targeted instances [78, 79]. Recent meta-analyses have demonstrated progress in the capacity to implement ML and differential abundance analyses for evaluation of multi-study cohorts via gradient boosting and random forest classifiers [80].

Training such models requires labeled “control” and “disease” observations; however, given the complex heterogeneity of microbiome study cohorts, it is important to carefully consider the handling of such data when evaluating multi-label assignments [81]. When control samples are included in the multi-label task, the model learns a global embedding space that encompasses both control and diseased microbiome states, potentially providing a reference baseline against which disease patterns can be contrasted. This approach allows the model to understand the full spectrum of microbiome variation, from control states through various disease conditions, which would help identify how different diseases represent deviations from baseline microbiome patterns. Their inclusion would enable the model to learn more generalizable features that capture the fundamental differences between health and disease, while simultaneously distinguishing between specific disease types within the embedding space. However, this approach faces the challenge that control samples cannot be meaningfully assigned disease type labels, creating supervision gaps where the model must learn to predict disease type and geographical/anatomical information while leaving the disease type undefined.

Excluding control samples from multi-label classification creates a specialized embedding space focused entirely on diseased microbiome states, allowing the model to concentrate on learning the subtle microbiome signatures that distinguish between different disease conditions. This approach enables the model to develop fine-grained representations of disease-specific microbiome patterns without the potentially overwhelming influence of controls that might dominate the embedding space and obscure important inter-disease differences. However, this approach sacrifices the potential benefits of having a control reference point, which might limit the model’s ability to understand how diseases represent specific deviations from normal microbiome states rather than simply learning arbitrary distinctions between disease patterns.

These considerations are critical to revealing important information regarding those microbiome associations that are consistent between disease states and those that are not, context that is critical to implementation of generalizable models.

### Embeddings to Evaluate Generalizability

The capacity to assess generalizability of our models is further supported by evaluating embedding spaces underlying the relevant classifiers. Embeddings can be used to project and evaluate data structure to assess the compatibility of a new, unseen dataset for inference via an existing model [65, 82, 83].

Our embedding space trained with control samples provides a framework for analyzing unseen microbiome samples for which disease state classification is desired by first establishing whether they represent control or diseased states, then progressively refining predictions about specific characteristics. This will facilitate future applications where high-level phenotypic classification is desired, followed by tailored multi-label predictions dependent on application-specific aims.

When an unknown sample is projected into this embedding space, its position relative to learned control and disease clusters could reveal its health status: samples clustering near controls suggest normal microbiomes with respect to their counterparts in the corresponding study), while those clustering with disease samples indicate disease-like states. This approach allows researchers to first answer the fundamental question of health versus disease, then drill down into specific disease types, body sites, and geographical origins for samples identified as diseased. The presence of controls in the embedding space provides crucial reference points that help calibrate the model’s confidence and establish meaningful distance metrics for determining how similar an unknown sample is to known control baselines. This is comparable to applications in other domains, where anomalous data points are identified via comparison to control data representing a baseline [84, 85, 86]. Our study-level hold-out analysis demonstrates that our approach is capable of classifying disease state for unseen, out-of-distribution datasets, and demonstrates potential for future assessments where out-of-study generalizability is desired.

In contrast, the disease-focused embedding space without controls offers superior discrimination for samples already suspected to be diseased, enabling finegrained classification of disease types. Such applications could be critically informative in instances of known off-normal clinical conditions for diseases with complex and heterogenous presentation, such as immune dysfunction [87]. Unknown samples projected into this specialized space would be compared directly against clusters of specific diseases, potentially revealing subtle pathological patterns that might be overlooked in a broader embedding space dominated by the large separation between control and diseased states. This approach excels in the event of a need to distinguish between similar diseases or identify rare pathological conditions, as the model has concentrated its representational capacity on capturing the differences between various diseases rather than the broader control-diseased distinction. A limitation is that an unseen normal sample might be misclassified as diseased since this version of the model lacks experience with normal patterns. Despite the potential for such limitations, the ability to evaluate and assign a preliminary classification label to a complex, idiopathic disease state would have tremendous value in environments where diagnostic practices are limited [88].

### Considerations for Future Implementation

The ability to assign clinically relevant classification labels to unseen microbiome specimens represents an important capability for advancing microbiome science to application, especially in the biomarker and diagnostic space [71]. The optimal strategy for analyzing unknown microbiomes might involve using multiple embedding spaces sequentially or in parallel to leverage their complementary strengths. Future efforts that further explore and integrate embeddings derived from multimodal data could improve generalizability for novel applications, including rare diseases without sufficient training data, an important clinical need [89].

The control-inclusive model described here could serve as an initial screening tool to determine overall health status and provide broad classification, while the disease-focused model could offer detailed pathological analysis for samples identified as potentially diseased. This dual approach would be particularly valuable in clinical settings where unknown samples could represent either control individuals requiring reassurance or patients with undiagnosed conditions requiring specific therapeutic interventions. Further supplementation of training data would strengthen each model. By comparing how unknown samples cluster in both embedding spaces, researchers could also identify novel microbiome states that do not fit well into either framework, potentially revealing new disease subtypes or unusual variants that expand our understanding of microbiome diversity.

## 5 Conclusions

We developed a fusion model trained on visual representations of microbiome abundance profiles and HFE features. By integrating ViT and CNN submodels, the model captures local and global features from microbiome map images. This enables the creation of embeddings that encode disease status, disease type, and geographical location. Our fusion model achieves high classification performance in differentiating disease states across a large metagenomic sample collection, and the generated embeddings form clusters that align with abundance distributions in disease states. The trained models can generate new embeddings for unseen samples that fit into existing clusters and can be used to label unknown samples.

Our collection of 13*K* image profiles represents, to our knowledge, the largest meta-study set for disease state prediction from metagenomic data, and the first to implement computer vision in such microbiome applications. This image collection includes curated labels for all samples, as well as clinical metadata [46].

Our trained fusion model can be used as a building block for follow-on machine learning applications that merge visual representations of abundance profiles with a multitude of other data modalities, and provides a strong foundation for creating models that enable prediction and characterization via the human gut microbiome in relevant disease states.

## Abbreviations

ViT: Vision Transformer
CNN: Convolutional Neural Network
HFE: Hierarchical Feature Engineering
HCV: Hilbert Curve Visualization
CLR: Centered Log Ratio
LOO: Leave One Out
BCE: Binary Cross Entropy
UMAP: Uniform Manifold Approximation and Projection
CoDA: Compositional Data Analysis
mWGS: Metagenomic Whole Genome Sequencing
16S: 16S Ribosomal RNA Sequencing
ASV: Amplicon Sequence Variant
HMP: Human Microbiome Project
LLNL: Lawrence Livermore National Laboratory

## Declarations

### Ethics Approval and Consent to Participate

Not Applicable.

### Consent for Publication

All Authors Reviewed the Manuscript and Consent to its Publication.

### Availability of Data and Material

All source code used for model training and sample inference can be found at the following GitHub repository: https://github.com/LLNL/invisorg. This study re-analyzed publicly available de-identified datasets from open literature. No new specimens were obtained or generated for this study, and the paper [46] contains all details and materials.

## Acknowledgements

The authors would like to acknowledge the multiple groups in the Biosciences & Biotechnology Division at Lawrence Livermore National Laboratory for their feedback and discussion. We also thank Dr. Jennifer Clarke at the University of Nebraska Lincoln for her feedback and comments on the results. Finally, we also would like to thank the high performance computing team at LLNL’s Livermore Computing group for their invaluable help with hardware and software computing resources.

## Additional Files

### Additional File 1 — Supplementary Materials

The supplementary materials document contains detailed descriptions of various aspects of our work, as well as supporting information regarding our analyses. The document contains the particulars of the datasets that we collected and used, as well as details regarding any preprocessing steps that were performed. Details on the database that was used for profiling the microbiome samples (and generating the abundance profiles), as well as model training specifics for our models can be found in this document.d

## References

1. Specter, M.: Germs are us. The New Yorker 88(Oct 22), 32–39 (2012)

2. Yong, E.: I contain multitudes: The microbes within us and a grander view of life 1 (2016)

3. Bordenstein, S.R., Theis, K.R.: Host biology in light of the microbiome: ten principles of holobionts and hologenomes. PLoS biology 13(8), 1002226 (2015)

4. Cho, I., Blaser, M.J.: The human microbiome: at the interface of health and disease. Nature Reviews Genetics 13(4), 260–270 (2012)

5. Jovel, J., Dieleman, L.A., Kao, D., Mason, A.L., Wine, E.: The human gut microbiome in health and disease. Metagenomics, 197–213 (2018)

6. Durack, J., Lynch, S.V.: The gut microbiome: Relationships with disease and opportunities for therapy. Journal of experimental medicine 216(1), 20–40 (2019)

7. Singh, B.K., Trivedi, P., Egidi, E., Macdonald, C.A., Delgado-Baquerizo, M.: Crop microbiome and sustainable agriculture. Nature Reviews Microbiology 18(11), 601–602 (2020)

8. Fierer, N.: Embracing the unknown: disentangling the complexities of the soil microbiome. Nature Reviews Microbiology 15(10), 579–590 (2017)

9. Swayambhu, M., Kümmerli, R., Arora, N.: Microbiome-based stain analyses in crime scenes. Applied and Environmental Microbiology 89(1), 01325–22 (2023)

10. Metcalf, J.L., Xu, Z.Z., Bouslimani, A., Dorrestein, P., Carter, D.O., Knight, R.: Microbiome tools for forensic science. Trends in biotechnology 35(9), 814–823 (2017)

11. Haitaian, C., Marchant, G.: Legal implications of the human microbiome. BUJ Sci. & Tech. L. 28, 39 (2022)

12. Minogue, T.D., Koehler, J.W., Stefan, C.P., Conrad, T.A.: Next-generation sequencing for biodefense: bio-threat detection, forensics, and the clinic. Clinical chemistry 65(3), 383–392 (2019)

13. Fazil, A.H.A., Reddy, U.S., Gumpu, M.B.: Biosensor for biothreat detection and defense application 1, 267–291 (2024)

14. Wooley, J.C., Godzik, A., Friedberg, I.: A primer on metagenomics. PLoS computational biology 6(2), 1000667 (2010)

15. Sleator, R.D., Shortall, C., Hill, C.: Metagenomics. Letters in applied microbiology 47(5), 361–366 (2008)

16. Kim, N., Ma, J., Kim, W., Kim, J., Belenky, P., Lee, I.: Genome-resolved metagenomics: a game changer for microbiome medicine. Experimental & Molecular Medicine 56(7), 1501–1512 (2024)

17. Kim, C., Pongpanich, M., Porntaveetus, T.: Unraveling metagenomics through long-read sequencing: A comprehensive review. Journal of Translational Medicine 22(1), 111 (2024)

18. Woodcroft, B.J., Aroney, S.T., Zhao, R., Cunningham, M., Mitchell, J.A., Blackall, L., Tyson, G.W.: Singlem and sandpiper: Robust microbial taxonomic profiles from metagenomic data. bioRxiv, 2024–01 (2024)

19. Majzoub, M.E., Luu, L.D., Haifer, C., Paramsothy, S., Borody, T.J., Leong, R.W., Thomas, T., Kaakoush, N.O.: Refining microbial community metabolic models derived from metagenomics using reference-based taxonomic profiling. mSystems, 00746–24 (2024)

20. Archana, T., Singh, S., Kumar, D., Kumar, V., Vaja, S., Kumar, G.: An insight into functional metagenomics profiling of different ecosystems, 417–430 (2024)

21. Sardar, P., Almeida, A., Pedicord, V.A.: Integrating functional metagenomics to decipher microbiome– immune interactions. Immunology and Cell Biology (2024)

22. Akintunde, O., Tucker, T., Carabetta, V.J.: The evolution of next-generation sequencing technologies, 3–29 (2024)

23. Jensen, T.B.N., Dall, S.M., Knutsson, S., Karst, S.M., Albertsen, M.: High-throughput dna extraction and cost-effective miniaturized metagenome and amplicon library preparation of soil samples for dna sequencing. Plos one 19(4), 0301446 (2024)

24. Yi, X., Lu, H., Liu, X., He, J., Li, B., Wang, Z., Zhao, Y., Zhang, X., Yu, X.: Unravelling the enigma of the human microbiome: Evolution and selection of sequencing technologies. Microbial Biotechnology 17(1), 14364 (2024)

25. Zhang, D., Li, X., Wang, Y., Zhao, Y., Zhang, H.: The clinical importance of metagenomic next-generation sequencing in detecting disease-causing microorganisms in cases of sepsis acquired in the community or hospital setting. Frontiers in Microbiology 15, 1384166 (2024)

26. Teixeira, M., Silva, F., Ferreira, R.M., Pereira, T., Figueiredo, C., Oliveira, H.P.: A review of machine learning methods for cancer characterization from microbiome data. NPJ Precision Oncology 8(1), 123 (2024)

27. Wu, J., Singleton, S.S., Bhuiyan, U., Krammer, L., Mazumder, R.: Multi-omics approaches to studying gastrointestinal microbiome in the context of precision medicine and machine learning. Frontiers in molecular biosciences 10, 1337373 (2024)

28. Sun, Z., Huang, S., Zhang, M., Zhu, Q., Haiminen, N., Carrieri, A.P., Vázquez-Baeza, Y., Parida, L., Kim, H.-C., Knight, R., et al.: Challenges in benchmarking metagenomic profilers. Nature methods 18(6), 618–626 (2021)

29. Fischer, M., Strauch, B., Renard, B.Y.: Abundance estimation and differential testing on strain level in metagenomics data. Bioinformatics 33(14), 124–132 (2017)

30. Nayfach, S., Pollard, K.S.: Toward accurate and quantitative comparative metagenomics. Cell 166(5), 1103–1116 (2016)

31. Yates, A.D., Allen, J., Amode, R.M., Azov, A.G., Barba, M., Becerra, A., Bhai, J., Campbell, L.I., Carbajo Martinez, M., Chakiachvili, M., et al.: Ensembl genomes 2022: an expanding genome resource for non-vertebrates. Nucleic acids research 50(D1), 996–1003 (2022)

32. Esteva, A., Robicquet, A., Ramsundar, B., Kuleshov, V., DePristo, M., Chou, K., Cui, C., Corrado, G., Thrun, S., Dean, J.: A guide to deep learning in healthcare. Nature medicine 25(1), 24–29 (2019)

33. Kussul, N., Lavreniuk, M., Skakun, S., Shelestov, A.: Deep learning classification of land cover and crop types using remote sensing data. IEEE Geoscience and Remote Sensing Letters 14(5), 778–782 (2017)

34. Hammoudeh, Z., Lowd, D.: Training data influence analysis and estimation: A survey. Machine Learning 113(5), 2351–2403 (2024)

35. Subramanian, S., Harrington, P., Keutzer, K., Bhimji, W., Morozov, D., Mahoney, M.W., Gholami, A.: Towards foundation models for scientific machine learning: Characterizing scaling and transfer behavior. Advances in Neural Information Processing Systems 36 (2024)

36. Jayakrishnan, T.T., Sangwan, N., Barot, S.V., Farha, N., Mariam, A., Xiang, S., Aucejo, F., Conces, M., Nair, K.G., Krishnamurthi, S.S., et al.: Multi-omics machine learning to study host-microbiome interactions in early-onset colorectal cancer. NPJ Precision Oncology 8(1), 146 (2024)

37. Roy, G., Prifti, E., Belda, E., Zucker, J.-D.: Deep learning methods in metagenomics: a review. Microbial Genomics 10(4), 001231 (2024)

38. Malakar, S., Sutaoney, P., Madhyastha, H., Shah, K., Chauhan, N.S., Banerjee, P.: Understanding gut microbiome-based machine learning platforms: A review on therapeutic approaches using deep learning. Chemical Biology & Drug Design 103(3), 14505 (2024)

39. Soueidan, H., Nikolski, M.: Machine learning for metagenomics: methods and tools. arXiv preprint arXiv:1510.06621 (2015)

40. Li, J., Xiong, A., Wang, J., Wu, X., Bai, L., Zhang, L., He, X., Li, G.: Deciphering the microbial landscape of lower respiratory tract infections: insights from metagenomics and machine learning. Frontiers in Cellular and Infection Microbiology 14, 1385562 (2024)

41. Lee, S., Lee, I.: Comprehensive assessment of machine learning methods for diagnosing gastrointestinal diseases through whole metagenome sequencing data. Gut Microbes 16(1), 2375679 (2024)

42. Yerke, A., Fry Brumit, D., Fodor, A.A.: Proportion-based normalizations outperform compositional data transformations in machine learning applications. Microbiome 12(1), 45 (2024)

43. Valdes, C., Stebliankin, V., Ruiz-Perez, D., Park, J.I., Lee, H., Narasimhan, G.: Microbiome maps: Hilbert curve visualizations of metagenomic profiles. Frontiers in Bioinformatics 3, 1154588 (2023)

44. Shang, J., Peng, C., Tang, X., Sun, Y.: Phavip: Phage virion protein classification based on chaos game representation and vision transformer. Bioinformatics 39(Supplement 1), 30–39 (2023)

45. Cadow, J., Manica, M., Mathis, R., Reddel, R.R., Robinson, P.J., Wild, P.J., Hains, P.G., Lucas, N., Zhong, Q., Guo, T., et al.: On the feasibility of deep learning applications using raw mass spectrometry data. Bioinformatics 37(Supplement 1), 245–253 (2021)

46. Kok, C.R., Mulakken, N., Thissen, J.B., Manuel Marti, J., Lee, R., Trainer, J.B., Goncalves, A., Ranganathan, H., Avila-Herrera, A., Jaing, C., et al.: Meta2db: Curated shotgun metagenomic feature sets and metadata for health state prediction. bioRxiv, 2024–10 (2024)

47. Kim, D., Song, L., Breitwieser, F.P., Salzberg, S.L.: Centrifuge: rapid and sensitive classification of metagenomic sequences. Genome research 26(12), 1721–1729 (2016)

48. Martí, J.M.: Recentrifuge: Robust comparative analysis and contamination removal for metagenomics. PLoS computational biology 15(4), 1006967 (2019)

49. Quinn, T.P., Erb, I., Gloor, G., Notredame, C., Richardson, M.F., Crowley, T.M.: A field guide for the compositional analysis of any-omics data. GigaScience 8(9), 107 (2019)

50. Greenacre, M.: Compositional data analysis. Annual Review of Statistics and its Application 8(1), 271–299 (2021)

51. Valdes, C.: Microbiome Maps. Software and documentation for Hilbert curve visualization of metagenomic profiles (2023). www.microbiomemaps.org/ Accessed 2024-12-19

52. Oudah, M., Henschel, A.: Taxonomy-aware feature engineering for microbiome classification. BMC bioinformatics 19, 1–13 (2018)

53. Kittler, J.: Combining classifiers: A theoretical framework. Pattern analysis and Applications 1, 18–27 (1998)

54. Morvant, E., Habrard, A., Ayache, S.: Majority vote of diverse classifiers for late fusion. In: Structural, Syntactic, and Statistical Pattern Recognition: Joint IAPR International Workshop, S+ SSPR 2014, Joensuu, Finland, August 20-22, 2014. Proceedings, pp. 153–162 (2014). Springer

55. Gadzicki, K., Khamsehashari, R., Zetzsche, C.: Early vs late fusion in multimodal convolutional neural networks. In: 2020 IEEE 23rd International Conference on Information Fusion (FUSION), pp. 1–6 (2020). IEEE

56. Dosovitskiy, A.: An image is worth 16×16 words: Transformers for image recognition at scale. arXiv preprint arXiv:2010.11929 (2020)

57. Vaswani, A., Shazeer, N., Parmar, N., Uszkoreit, J., Jones, L., Gomez, A.N., Kaiser, L., Polosukhin, I.: Attention is all you need.(nips), 2017. arXiv preprint arXiv:1706.03762 10, 0140525–16001837 (2017)

58. LeCun, Y., Boser, B.E., Denker, J.S., Henderson, D., Howard, R.E., Hubbard, W.E., Jackel, L.D.: Back-propagation Applied to Handwritten Zip Code Recognition. Neural Computation 1(4), 541–551 (1989). doi:10.1162/neco.1989.1.4.541

59. Lu, K., Xu, Y., Yang, Y.: Comparison of the potential between transformer and cnn in image classification. In: ICMLCA 2021; 2nd International Conference on Machine Learning and Computer Application, pp. 1–6 (2021). VDE

60. Li, Z., Liu, F., Yang, W., Peng, S., Zhou, J.: A survey of convolutional neural networks: analysis, applications, and prospects. IEEE transactions on neural networks and learning systems 33(12), 6999–7019 (2021)

61. Pulfer, B.: Vision Transformers from Scratch PyTorch: A Step-by-Step Guide. https://medium.com/@brianpulfer. Accessed: May 1, 2024 (2022)

62. Vilas, M.G., Schaumlöffel, T., Roig, G.: Analyzing vision transformers for image classification in class embedding space. Advances in neural information processing systems 36 (2024)

63. Truong, T., Jush, F.K., Lenga, M.: Benchmarking pretrained vision embeddings for near-and duplicate detection in medical images. In: 2024 IEEE International Symposium on Biomedical Imaging (ISBI), pp. 1–5 (2024). IEEE

64. Consortium, U.: Uniprot: the universal protein knowledgebase in 2023. Nucleic acids research 51(D1), 523–531 (2023)

65. Odrzywolek, K., Karwowska, Z., Majta, J., Byrski, A., Milanowska-Zabel, K., Kosciolek, T.: Deep embeddings to comprehend and visualize microbiome protein space. Scientific Reports 12(1), 10332 (2022)

66. Contributors, T.: Torchvision: Models, Datasets and Transformations for Images. https://github.com/pytorch/vision (2016)

67. Deng, J., Dong, W., Socher, R., Li, L.-J., Li, K., Fei-Fei, L.: Imagenet: A large-scale hierarchical image database. In: 2009 IEEE Conference on Computer Vision and Pattern Recognition, pp. 248–255 (2009). Ieee

68. Ferreiro, A.L., Choi, J., Ryou, J., Newcomer, E.P., Thompson, R., Bollinger, R.M., Hall-Moore, C., Ndao, I.M., Sax, L., Benzinger, T.L., et al.: Gut microbiome composition may be an indicator of preclinical alzheimer’s disease. Science translational medicine 15(700), 2984 (2023)

69. Le Goallec, A., Tierney, B.T., Luber, J.M., Cofer, E.M., Kostic, A.D., Patel, C.J.: A systematic machine learning and data type comparison yields metagenomic predictors of infant age, sex, breastfeeding, antibiotic usage, country of origin, and delivery type. PLoS computational biology 16(5), 1007895 (2020)

70. Fukui, H., Nishida, A., Matsuda, S., Kira, F., Watanabe, S., Kuriyama, M., Kawakami, K., Aikawa, Y., Oda, N., Arai, K., et al.: Usefulness of machine learning-based gut microbiome analysis for identifying patients with irritable bowels syndrome. Journal of clinical medicine 9(8), 2403 (2020)

71. Kumar, B., Lorusso, E., Fosso, B., Pesole, G.: A comprehensive overview of microbiome data in the light of machine learning applications: categorization, accessibility, and future directions. Frontiers in Microbiology 15, 1343572 (2024)

72. Arif, S.J., Graham, S.P., Abdill, R.J., Blekhman, R.: Analyzing human gut microbiome data from global populations: challenges and resources. Trends in Microbiology (2025)

73. Nearing, J.T., Douglas, G.M., Hayes, M.G., MacDonald, J., Desai, D.K., Allward, N., Jones, C.M., Wright, R.J., Dhanani, A.S., Comeau, A.M., et al.: Microbiome differential abundance methods produce different results across 38 datasets. Nature communications 13(1), 342 (2022)

74. Soenksen, L.R., Ma, Y., Zeng, C., Boussioux, L., Villalobos Carballo, K., Na, L., Wiberg, H.M., Li, M.L., Fuentes, I., Bertsimas, D.: Integrated multimodal artificial intelligence framework for healthcare applications. NPJ digital medicine 5(1), 149 (2022)

75. Nguyen, P.T., Di Rocco, J., Iovino, L., Di Ruscio, D., Pierantonio, A.: Evaluation of a machine learning classifier for metamodels. Software and Systems Modeling 20(6), 1797–1821 (2021)

76. Feurer, M., Klein, A., Eggensperger, K., Springenberg, J., Blum, M., Hutter, F.: Efficient and robust automated machine learning. Advances in neural information processing systems 28 (2015)

77. Liu, Y., Fachrul, M., Inouye, M., Méric, G.: Harnessing human microbiomes for disease prediction. Trends in microbiology 32(7), 707–719 (2024)

78. Jiang, C., Yang, J., Peng, X., Li, X.: A permutable mlp-like architecture for disease prediction from gut metagenomic data. BMC bioinformatics 25(1), 246 (2024)

79. Tang, J.-W., Tay, A.C.Y., Wang, L.: Interpretive prediction of hyperuricemia and gout patients via machine learning analysis of human gut microbiome. BMC microbiology 25(1), 429 (2025)

80. Jin, D.-M., Morton, J.T., Bonneau, R.: Meta-analysis of the human gut microbiome uncovers shared and distinct microbial signatures between diseases. Msystems 9(8), 00295–24 (2024)

81. Singla, K., Biswas, S.: Machine learning explanability method for the multi-label classification model. In: 2021 IEEE 15th International Conference on Semantic Computing (ICSC), pp. 337–340 (2021). IEEE

82. Pope, Q., Varma, R., Tataru, C., David, M.M., Fern, X.: Learning a deep language model for microbiomes: the power of large scale unlabeled microbiome data. PLOS Computational Biology 21(5), 1011353 (2025)

83. Oh, M., Zhang, L.: Deepmicro: deep representation learning for disease prediction based on microbiome data. Scientific reports 10(1), 6026 (2020)

84. Dörr, A.-K., Imangaliyev, S., Karadeniz, U., Schmidt, T., Meyer, F., Kraiselburd, I.: Distinguishing critical microbial community shifts from normal temporal variability in human and environmental ecosystems. Scientific Reports 15(1), 16934 (2025)

85. Gerstung, M., Pellagatti, A., Malcovati, L., Giagounidis, A., Porta, M.G.D., Jädersten, M., Dolatshad, H., Verma, A., Cross, N.C., Vyas, P., et al.: Combining gene mutation with gene expression data improves outcome prediction in myelodysplastic syndromes. Nature communications 6(1), 5901 (2015)

86. Xu, K., Wu, Q., Lu, Y., Zheng, Y., Li, W., Tang, X., Wang, J., Sun, X.: Meatrd: Multimodal anomalous tissue region detection enhanced with spatial transcriptomics. In: Proceedings of the AAAI Conference on Artificial Intelligence, vol. 39, pp. 12918–12926 (2025)

87. Ruscitti, P., Cantarini, L., Ciccia, F., Conti, F., Dagna, L., Iannone, F., Montecucco, C., Giovanni, P., Sfriso, P., Giacomelli, R.: Managing the clinical heterogeneity of patients with still’s disease, from early diagnosis to timely treatment. Autoimmunity Reviews, 103880 (2025)

88. Song, Y., Kono, M.: Potential biomarkers in systemic lupus erythematosus. JMA journal 8(3), 689–698 (2025)

89. Souza, R., Mouches, P., Wilms, M., Tuladhar, A., Langner, S., Forkert, N.D.: An analysis of the effects of limited training data in distributed learning scenarios for brain age prediction. Journal of the American Medical Informatics Association 30(1), 112–119 (2022)

